# Analyzing dynamic decision-making models using Chapman-Kolmogorov equations

**DOI:** 10.1101/588087

**Authors:** Nicholas W. Barendregt, Krešimir Josić, Zachary P. Kilpatrick

**Author notes:** K. Josić and Z.P. Kilpatrick share equal authorship.

## Abstract

Decision-making in dynamic environments typically requires adaptive evidence accumulation that weights new evidence more heavily than old observations. Recent experimental studies of dynamic decision tasks require subjects to make decisions for which the correct choice switches stochastically throughout a single trial. In such cases, an ideal observer’s belief is described by an evolution equation that is doubly stochastic, reflecting stochasticity in the both observations and environmental changes. In these contexts, we show that the probability density of the belief can be represented using differential Chapman-Kolmogorov equations, allowing efficient computation of ensemble statistics. This allows us to reliably compare normative models to near-normative approximations using, as model performance metrics, decision response accuracy and Kullback-Leibler divergence of the belief distributions. Such belief distributions could be obtained empirically from subjects by asking them to report their decision confidence. We also study how response accuracy is affected by additional internal noise, showing optimality requires longer integration timescales as more noise is added. Lastly, we demonstrate that our method can be applied to tasks in which evidence arrives in a discrete, pulsatile fashion, rather than continuously.

## 1 Introduction

Natural environments are fluid, and living beings need to accumulate evidence adaptively in order to make sound decisions (Behrens et al., 2007; Ossmy et al., 2013). Theoretical models suggest, and experiments confirm, that in changing environments animals use decision strategies that value recent observations more than older ones (Yu and Cohen, 2008; Brea et al., 2014; Urai et al., 2017). For instance, adaptive evidence accumulation has been explored using a dynamic version of the random dot motion discrimination (RDMD) task (Glaze et al., 2015). In this task, subjects must determine the predominant direction (left or right) of a field of randomly moving dots while this direction switches stochastically according to a continuous time Markov process. Since switches are unpredictable, an ideal observer discounts old information in favor of new evidence. Furthermore, this discounting rate increases with the rate of environmental changes. This strategy has been observed in humans and other animals performing dynamic tasks (Glaze et al., 2015; Piet et al., 2018; Glaze et al., 2018).

Normative models and their approximations have been used successfully to understand how subjects make decisions (Ratcliff, 1978; Gold and Shadlen, 2007). In simple cases these models are tractable and make concrete predictions about response statistics that can be compared to experimental data (Bogacz et al., 2006; Drugowitsch, 2016; Ratcliff and McKoon, 2008). However, determining when subjects use approximately normative decision strategies, and when and how they fail to do so, can be computationally challenging. For instance, one may wish to study how a subject’s estimate of the environmental timescale impacts their response accuracy, or how heuristic evidence-discounting strategies compare to optimal ones (Glaze et al., 2018; Radillo et al., 2019). To address these questions, previous work has primarily relied on Monte Carlo simulations (Veliz-Cuba et al., 2016; Piet et al., 2018), which can be computationally expensive.

Here, we show how to reframe dynamic decision models by deriving corresponding differential Chapman-Kolmogorov (CK) equations (See Eq. (6)). This approach allows us to quickly compute observer beliefs and performance, and compare models. Realizations of our models are described by stochastic differential equations with a drift term that switches according to a two-state Markov process, and leak terms that discount evidence. To describe these models using CK equations, we treat the switching process as a source of dichotomous noise, and condition on its state to track conditional belief densities. These methods allow us to quickly answer questions about how characteristics of optimal models and their approximations vary across ranges of task parameters.

Nonlinear, normative models can thus be compared to approximate linear and cubic discounting models, models with internal noise, and explicitly solvable bounded accumulation models with no flux boundaries. These models all can obtain near-optimal response accuracy, but each has very different belief distributions. This suggests that subject confidence reports could be used to distinguish subject decision strategies in data.

Detailed analyses, including belief distribution calculations, can be performed rapidly and accurately with our methods, allowing us to see *why* each approximate model performs better at different task difficulty levels. Monte Carlo methods fare much worse in terms of computation time and accuracy (See Fig. 9). Our methods also extend to tasks with pulsatile evidence, where drift and diffusion are replaced by jump terms. Our work thus demonstrates how partial differential equation descriptions of stochastic decision models, previously successful in understanding decision making in static environments (Busemeyer and Townsend, 1992; Moehlis et al., 2004; Bogacz et al., 2006), can be extended to dynamic environments.

## 2 Normative models for dynamic decision-making

We begin by considering the dynamic RDMD task (Glaze et al., 2015; Veliz-Cuba et al., 2016); an observer looks at a screen of dots which move, on average, right or left. The average direction of motion, which we call the state *s*(*t*), switches in time between states *s*_+_ (right-moving) and *s*_−_ (left-moving) as a two-state continuous time Markov process with hazard rate *h*, so *P* (*s*(*t* + Δ*t*) ≠ *s*(*t*)) = *h*Δ*t* + *o*(Δ*t*). The observer is interrogated at a random time, *T*, and reports their belief about the current direction of motion, *s*(*T*). The most reliable state estimate is obtained by computing the log-likelihood ratio (LLR) between choices from (noisy) observations, ξ(*t*), of the moving dot stimulus. Assuming the observer maintains a fixed estimate of the environmental hazard rate, 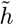, this evidence-accumulation process converges to a single stochastic differential equation (SDE) for the belief *y*(*t*) of the observer (See Veliz-Cuba et al. (2016) and Appendix A for modeling assumptions and details):

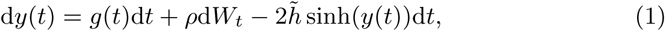

where *g*(*t*) is a telegraph process that switches between two values, ±*g*, with transition rate *h*, providing evidence about the state, *s*(*t*), d*W*_t_ is an increment of a Wiener process scaled by *ρ*, and the observer’s assumed hazard rate, 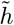, shapes the evidence discounting process. If we assume observations of the state *s*(*t*) are drawn from normal distributions, the input to the evidence accumulation model can be described by a single parameter (Veliz-Cuba et al., 2016). Combining our assumptions and rescaling time as *ht* ↦ *t*, we obtain the following SDE for the observer’s belief (See Appendix A) in rescaled time (different from the units in Eq. (1)):

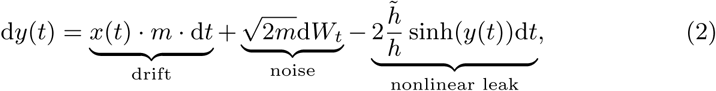

where *x*(*t*) ∈ ±1 is a telegraph process with switching rate equal to 1. The parameter *m* gives the mean information gain of the observer over the average length of time the environment remains the same (*h*^−1^ in original units, 1 in rescaled units). As *m* increases the task becomes easier. Thus, we refer to *m* as the *evidence strength*. If we take 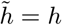, the explicit dependence of Eq. (2) on *h* vanishes, and, as we show, the observer obtains maximal response accuracy.

We are primarily interested in how variations of the evidence strength, *m*, true hazard rate, *h*, and the observer’s hazard rate estimate, 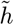, impact the response accuracy of an observer whose belief is represented by Eq. (2). These quantities can be changed by varying psychophysical task parameters (Glaze et al., 2015; Piet et al., 2018; Glaze et al., 2018), and so provide a means of validating Eq. (2) and its approximations. In addition, a thorough understanding of the normative model’s performance can provide insights into task parameter ranges in which a subject’s belief, *y*(*t*), is sensitive to the strategy they use (Radillo et al., 2019). Obtaining statistics of the solutions to Eq. (2) requires estimating the distribution of the stochastically evolving belief *y*(*t*) across time. Monte Carlo approaches can require many realizations to accurately characterize belief distributions (See Fig. 9 in Appendix B), and can thus be computationally prohibitive.

### 2.1 Expressing models using differential Chapman Kolmogorov equations

An alternative to sampling is to derive differential CK equations corresponding to Eq. (2) and evolve them to obtain time-dependent probability distributions, *p*(*y, t*), of observer belief, *y*(*t*), directly. For instance, for a fixed realization of *x*(*t*), the evolution of *p*(*y, t*) is described by the following differential CK equation:

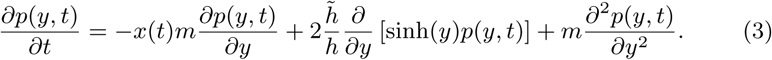

Here the drift terms involve non-autonomous forcing by *x*(*t*) and evidence discounting, while the diffusion term arises from the Wiener process. This equation could be useful for model fitting, since an experimenter would know the realization of *x*(*t*), and could then fit the single free parameter 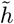 using response data. A simulation using a fixed realization of *x*(*t*) shown in Fig. 1A, reveals how the belief density tracks the state changes, and the peak of the distribution tends towards the fixed points 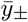 of Eq. (2) where 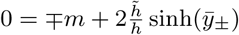.

**Fig. 1.**
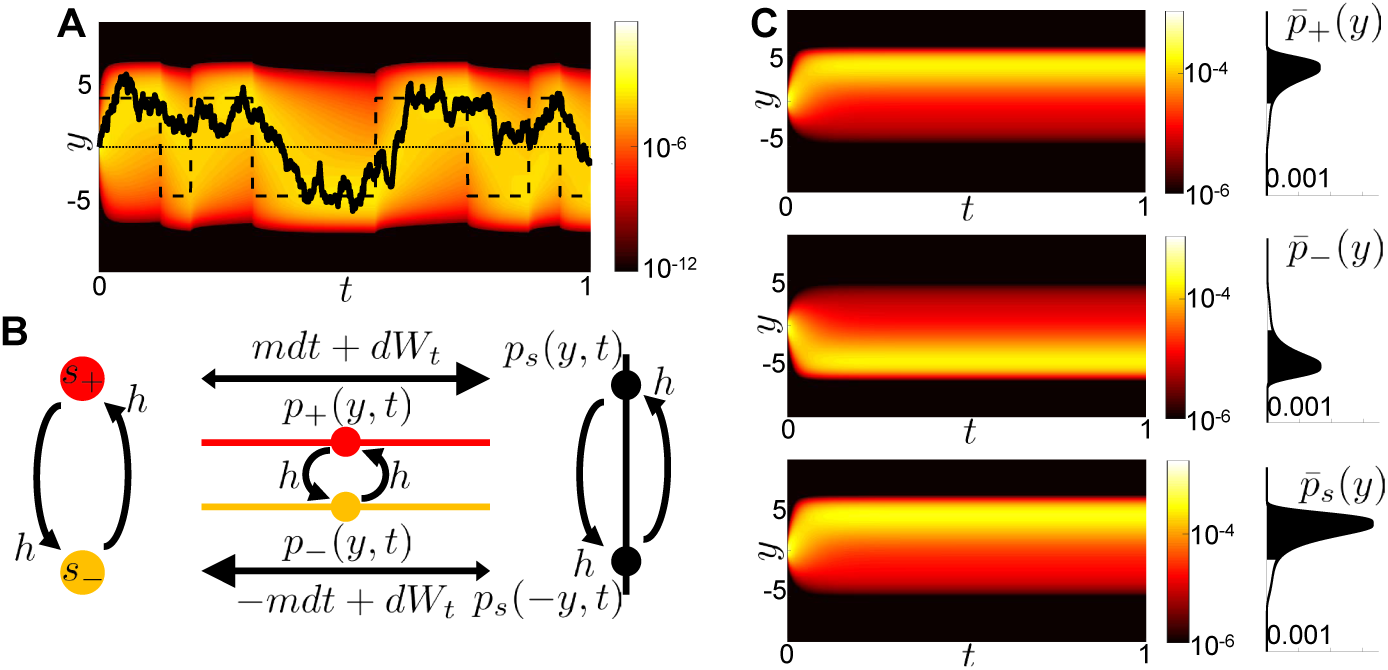
The evolution of solutions to the differential Chapman-Kolmogorov (CK) equations. **A:** Evolution of the probability density given by Eq. (3) for a fixed realization of the telegraph process, *x*(*t*). A single realization of the belief, *y,* (solid) and the instantaneous fixed point of Eq. (2) in the absence of Wiener process noise (dashed) are superimposed. **B:** Schematic of the differential CK equations. **C:** Sample evolution of densities *p*±(*y, t*) and *p*_*s*_(*y, t*) using Eqs. (4), and (6) respectively. Shaded region shows probability that contributes to accuracy of observer, computed using Eq. (5). Details on numerical methods are provided in Appendix B.

The evolution of the belief and performance across trials is determined by extending our model to include the distribution of possible realizations of *x*(*t*). Treating *x*(*t*) as dichotomous noise and defining the joint probability densities *p*±(*y, t*) := *p*(*y, t*|*s*(*t*) = *s*±), we obtain a set of coupled differential CK equations (Gardiner, 2004):

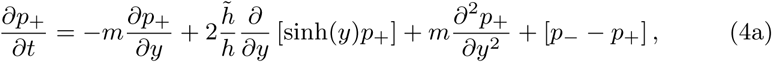

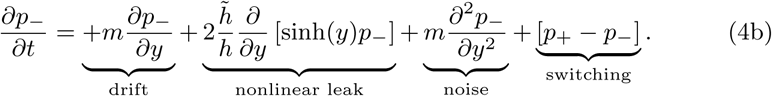

The jump terms that exchange probability between *p*_+_ and *p*_−_ in Eqs. (4) arise from the switches in state, as schematized in Fig. 1B. Eqs. (4) describe the joint evolution of the density of beliefs across all realizations of *x*(*t*). We assume symmetric priors, 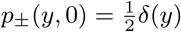. As we will show, this and the symmetry of Eq. (4) leads to symmetric solutions *p*_+_(*y, t*) = *p*_−_(−*y, t*).

Response accuracy – the probability of a correct response – is a common measure of subject performance in decision making tasks (Gold and Shadlen, 2007; Ratcliff and McKoon, 2008). Experimentally, response accuracy is defined as the fraction of correct responses at a specific interrogation time *T* (Glaze et al., 2015; Piet et al., 2018). In our model, optimal observers make choices in accordance with the sign of their belief, sign[*y*(*t*)], and response accuracy can be computed from solutions to Eq. (4) by computing

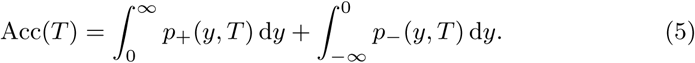

The belief, *y*(*t*), is *correct* if it has the same sign as *s*(*t*). This fact along with the inherent odd symmetry of Eqs. (4) suggests a change of variables −*y*↦ *y* in Eq. (4b). The sum, *p*_*s*_(*y, t*) := *p*_+_(*y, t*) + *p*_−_(−*y, t*), then evolves according to

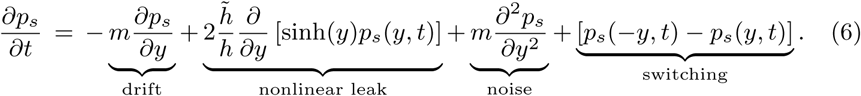

The new density *p*_*s*_(*y, t*) defines all belief values *y* > 0 as correct, since the sign of beliefs in the state *s*_−_ have been flipped, −*y* ↦ *y*, while the sign of all beliefs in state *s*_+_ remain the same. The density *p*_*s*_(*y, t*) thus describes beliefs *relative* to the state, *s*(*t*), with each environmental change flipping the sign of the belief, *y*(*t*) (See Fig. 1B). Eq. (5) can therefore be rewritten more simply as 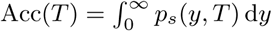. By symmetry, we can recover the two original densities as 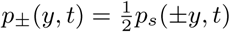.

Solving the CK equations numerically, we observe several notable features of *p*±(*y, t*) and *p*_*s*_(*y, t*) (Fig. 1C). First, the densities *p*±(*y, t*) are reflections of one another (*p*_+_(*y, t*) = *p*_−_(−*y, t*)) due to the symmetry of Eqs. (4). Second, all densities obtain stationarity on the timescale *h*^−1^ of the environment, so each is a unimodal function peaked on the *correct* side of *y* = 0. Stationary is reached due to the eventual equilibration between the drift and state switching. Most of the mass of the stationary densities is on the correct side of *y* = 0, and Acc(*T*) > 0.5. The long tail of the distribution *p*_*s*_(*y, t*) is due to both the constant transfer of probability from *y* to −*y* due to the switching and the Wiener process noise. Both the nonlinear leak and switching cause the accuracy Acc(*T*) to saturate over time.

Before going further, we note that Eq. (6) satisfies the conditions for existence of an ergodic process (Gardiner, 2004): The nonzero jump probabilities, and a positive diffusion coefficient, ensure that the differential CK equation converges to a unique stationary density as *t* → ∞. This occurs in a relatively short time period; we therefore focus the remainder of our study on steady-state cases. Typically, experimental dynamic decision trials are sufficiently long to make this assumption of stationarity reasonable (Glaze et al., 2015; Piet et al., 2018).

### 2.2 Evaluating accuracy for mistuned evidence-discounting

Subjects performing decision tasks often must learn the task parameters online to improve their performance. Our model can be extended to consider hazard rate learning (Radillo et al., 2017; Glaze et al., 2018), but for now we assume that the observer uses a fixed estimate 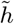 of the hazard rate for their evidence discounting strategy (Glaze et al., 2015).

How does the response accuracy of an observer whose belief is described by Eq. (2) change when 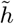 is mistuned? Veliz-Cuba et al. (2016) addressed this question using Monte Carlo sampling, but computational costs prevented a complete answer. Since Eq. (2) is rescaled, we take *h* = 1 for the remainder of our investigation; all other cases can be recovered by rescaling time. Before asking how changing 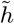 alters accuracy, we first briefly mention how accuracy varies with *evidence strength*, fixing 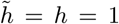. The density *p*_*s*_(*y, t*) computed using Eq. (6) rapidly converges to the stationary solution, with most of its mass above zero (Fig. 2A). As *m*, increases, more mass of the stationary distribution moves to positive values (Fig. 2B), but the total mass, equal to lim_*T*→∞_ Acc(*T*), always saturates at a value less than 1 due to discounting and state switching.

**Fig. 2.**
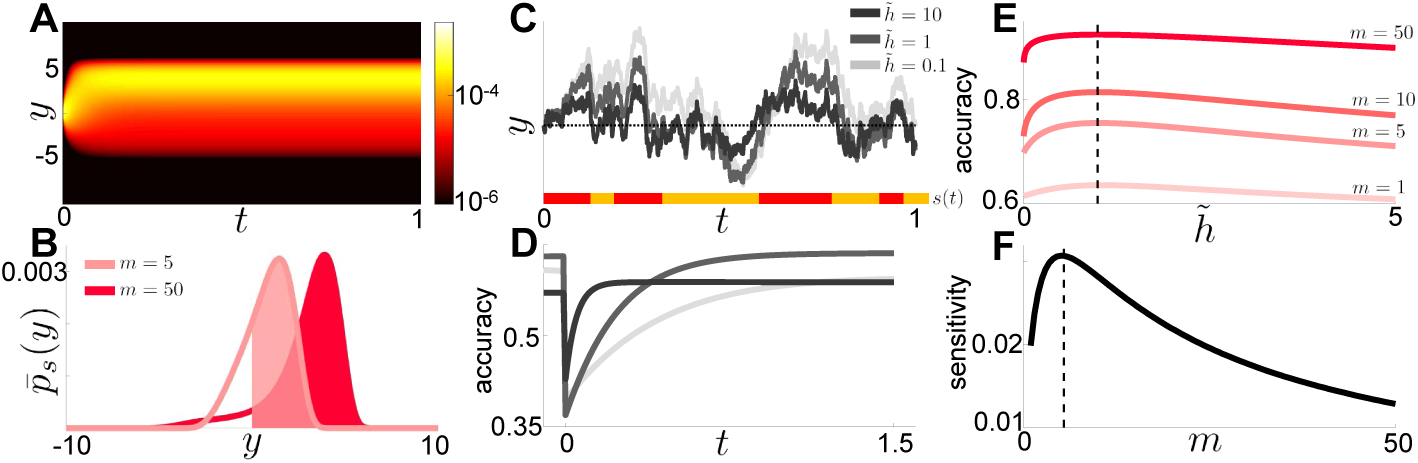
Performance of normative and mistuned nonlinear observer models. **A:** Evolution of the density *ps*(*y, t*) for *m* = 50 given by Eq. (6). **B:** Stationary densities, 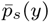, for *m* = 5, 50. With stronger evidence (*m* = 50), the stationary distribution has more mass above *y* = 0. **C:** Realizations of the belief variable *y* for different estimated hazard rates 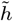 and fixed true hazard rate, *h* = 1. The environmental state, *s*(*t*), is shown below (red: *s*_+_; yellow: *s*_−_). **D:** Observer accuracy, for varied 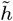, following a change point at *t* = 0. **E:** Observer steady-state accuracy as a function of 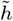 for different values of *m* is maximized when 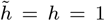. **F:** Curvature of the accuracy functions in **E** at the maximum, *h* = 1, as a function of *m*, shows the observer is most sensitive to changes in their estimated hazard rate 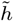 when *m* ≈ 5.

When the observer misestimates the hazard rate, 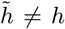, we expect the long term accuracy to suffer. Effects on accuracy are subtle, but do follow a general pattern: overestimating the hazard rate 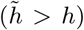 causes the observer to discount prior evidence too strongly, resulting in more errors driven by observation noise (Fig. 2C). On the other hand, observers that underestimate the hazard rate 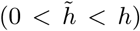 discount evidence too slowly and are less adaptive to change points. Change point triggered response accuracy plots show both of these trends (Fig. 2D). Accuracy obtains a lower ceiling value during longer epochs without environmental changes when the discounting rate 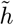 is too high. On the other hand, accuracy recovers more slowly following changes when the discounting rate 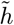 is too low. This bias-variance tradeoff is common to binary choice experiments in dynamic environments (Glaze et al., 2015, 2018): Low discounting rates lead to averaging over longer sequences of observations thus reducing the effect of observational noise while increasing bias. On the other hand, high discounting rates decrease bias but increase susceptibility to observational noise, resulting in higher variability. An optimal observer balances these two sources of inaccuracy at a given environmental hazard rate.

An experimenter may not be able to change discounting rate a subject uses, but can control the strength of evidence the subject integrates. We therefore asked how the accuracy of both ideal and mistuned observers is impacted by changes in *m*. Not surprisingly, accuracy increases as the strength of evidence increases (Fig. 2E). More interestingly, the sensitivity (or curvature) of the accuracy function at the optimum, where 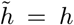, varies nonmonotonically with *m*, obtaining a peak at *m* ≈ 5 (Fig. 2F). Thus response accuracy is most sensitive to model mistuning for tasks of intermediate difficulty. Intuitively, an observer will always perform close to chance (Acc ≈ 0.5) when the task is hard (*m* is low), regardless of the *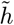* they use. The observer will perform well (Acc ≈ 1), again regardless of 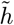, if the task is easy (*m* is high). At intermediate values of *m*, the observer’s performance is sensitive to changes in the model. See Radillo et al. (2019) for a similar analysis for a dynamic decisions using pulsatile evidence.

This example illustrates how CK equations can be used to obtain response accuracy statistics, and to compare normative models to related nonlinear models in which the evidence discounting is mistuned. Such approximate models may offer plausible descriptions of subject’s strategies, but only capture some of the possibilities. In the next section, we develop and analyze linear discounting models that approximate the adaptive evidence accumulation properties of the normative model and can be tuned to obtain near-optimal response accuracy.

## 3. Linear evidence discounting in dynamic environments

The nonlinear model defined by Eq. (2) describes the optimal evidence-accumulation strategy when the estimated hazard rate is correct. However, approximate models can also obtain response accuracy that is near-optimal. Glaze et al. (2015) and Veliz-Cuba et al. (2016) demonstrated this using a model that includes a linear leak term, *-λy,* in place of the nonlinearity in the normative model. The linear model is more tractable and can capture the dynamics of subjects’ beliefs in behavioral data (Ossmy et al., 2013; Glaze et al., 2015; Piet et al., 2018). We are interested in how well its statistics can be matched to that of the nonlinear model and how sensitive this match is to perturbations in the leak rate.

The linear discounting model is the doubly stochastic differential equation,

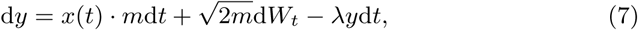

where *λ* is a parameter we tune. As before, we can write differential CK equations corresponding to Eq. (7), and define *p*_*s*_(*y, t*) = *p*_+_(*y, t*) + *p*_−_(−*y, t*) to obtain the evolution equation

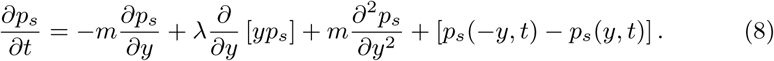

As in the nonlinear model, an attracting stationary solution to Eq. (8) exists as long as *λ* > 0 (Gardiner, 2004). We thus focus on stationary solutions, 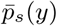, and make comparisons with the normative model. Our goal is to see how the leak rate, *λ,* can be tuned so that the behavior of an observer whose belief evolves according tof Eq. (7) best matches that of an observer using the normative model, Eq. (2).

We use two metrics to compare our models: first, we consider the accuracy of the linear model

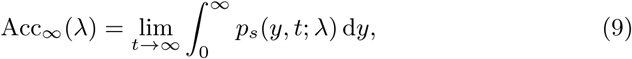

and aim to tune *λ* so Acc_∞_(*λ*) is maximized. Second, to quantify the distance between the belief distributions, we compute the Kullback-Leibler (KL) divergence

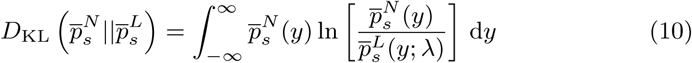

between the stationary normative distribution, 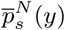, obtained from Eq. (6), and the stationary distribution of the linear approximation, 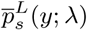, obtained from Eq. (8). While it is possible for models to have nearby belief distributions but different realizations within trials, minimizing KL divergence still penalizes models with divergent belief distributions, sure to have distinct trial wise realizations. We show that choosing the leak rate, *λ*, that maximizes accuracy or minimizes the KL divergence leads to different biases (Fig. 3A,B).

**Fig. 3.**
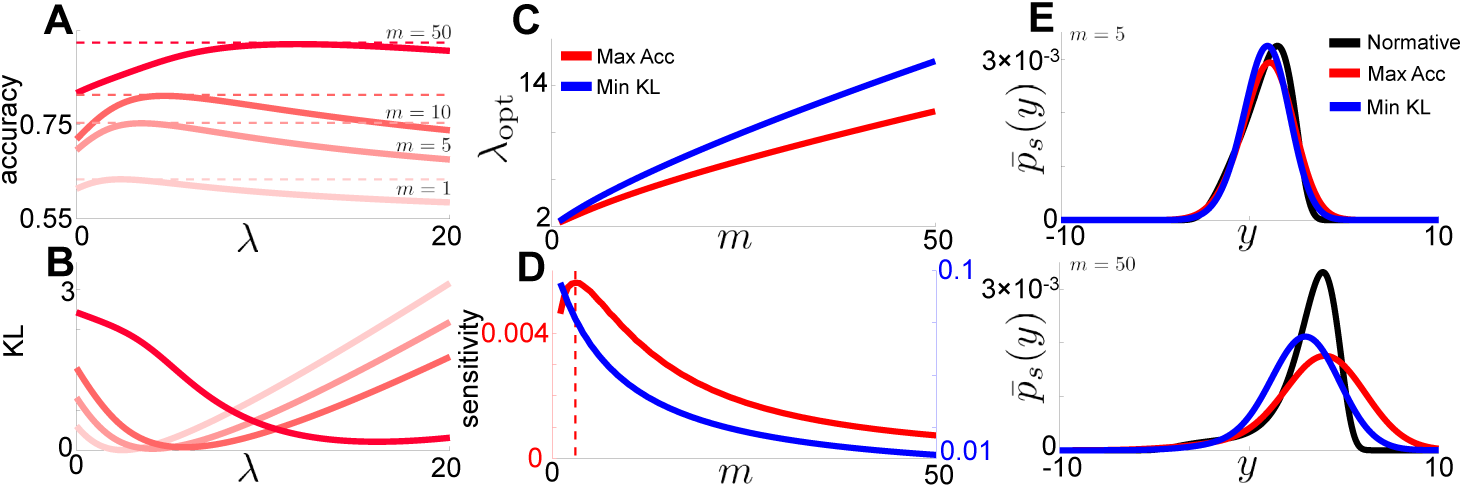
Linear two-alternative forced-choice task model. **A:** Performance of the linear observer as a function of discounting rate, *λ,* for different evidence strengths, *m*. **B:** KL divergence between the densities evolving under the normative and linear model as a function of *λ,* for different values of *m*. **C:** Optimal discounting rate for each metric as a function of *m*. **D:** Sensitivity of discounting rate on optimality metric as a function of *m*. **E:** Comparison of the normative density, and the linear density with 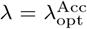 chosen to maximize accuracy (red), or with 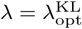 chosen to minimize KL divergence (blue) for *m* = 5 (top) and *m* = 50 (bottom).

Similar to the nonlinear model, the response accuracy of an observer using linear discounting varies nonmonotonically with *λ* (Fig. 3A). Observers using small *λ* adapt too slowly to change points, and those using a high *λ* exhibit more noise-driven errors in the state estimate. The optimal value of *λ* is achieved by balancing these error sources, obtaining response accuracy levels very close to those of the normative model. Furthermore, the 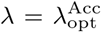 that maximizes response accuracy increases as *m* is increased, since evidence needs to be discounted more rapidly in environments with higher evidence strengths (Fig. 3C). The KL divergence also varies nonmonotonically with *λ* for all values of *m* (Fig. 3B), obtaining a minimum at a value 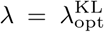 that also increases with *m*, but is higher than 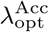. Understanding this result requires a more detailed analysis of the stationary densities of the normative and linear models, as we discuss below.

How important is it to tune *λ* in the linear model? If linear models are more sensitive to fine tuning for some task parameter ranges than the normative model, experimentalists could use these task parameter ranges to distinguish subjects’ strategies. When *m* is small the belief distributions 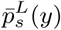 and 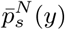 are close whether 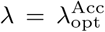, or 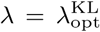 (Fig. 3C,E), but this agreement is sensitive to changes in *λ* (Fig. 3D). Thus, both the accuracy and KL divergence are sensitive to *λ* when *m* is small. For large *m* the two belief distributions are not close (Fig. 3E), and the KL divergence and difference in accuracy are insensitive to changes in *λ*. This disagreement in belief distribution at high *m* is less important when optimizing the accuracy of the linear model, as we only need to maximize the mass of 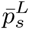 above *y* = 0.

Differences between the two models become apparent if we interrogate observers about their confidence and not just their choice (Fig. 3E). We hypothesize that if one were to fit behavioral data using response accuracy, and compare them to fits using subject’s confidence reports, the second approach would result in stronger leak rates. Indeed, considerations of subject confidence as a proxy for LLR has been an important development in recent decision making studies (Kiani and Shadlen, 2009; Van Den Berg et al., 2016), and we will revisit this view in Section 7. However, most behavioral studies of decisions in dynamic environments do not include confidence reports, and so model fits are typically performed by considering accuracy data. We thus focus primarily on this measure for the remainder of our study.

## 4 Tuning evidence accumulation to account for internal noise

We next explore the impact of additional noise sources on the performance of both the nonlinear and linear models. Since the nervous system is inherently noisy (Faisal et al., 2008), it is important to consider sources of variability on top of the stochasticity of observations when developing and fitting decision models (Smith, 2010). Brunton et al. (2013) showed that the responses of humans and rats in an auditory clicks task are best described by models that include internally generated noise. Piet et al. (2018) showed that the same is the case in a dynamic clicks task. With this in mind, we extend our analysis to incorporate an additional independent noise source. Such variability could arise in early sensory areas or as part of the decision process (Bankó et al., 2011). For simplicity, we model the source of noise as an independent Wiener process, *X*_*t*_, with variance scaled by a parameter *D*. The nonlinear model then takes the form,

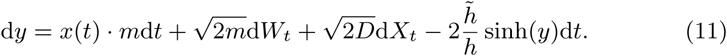

Adding internal noise means that Eq. (11) is no longer a normative model: When *D* > 0, noise corrupts state estimates (Fig. 4A), and maximal response accuracy is achieved when 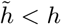, as we show. The linear model is updated similarly,

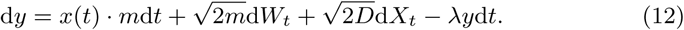

In either model, the Wiener processes, d*W*_*t*_ and d*X*_*t*_, are independent.

**Fig. 4.**
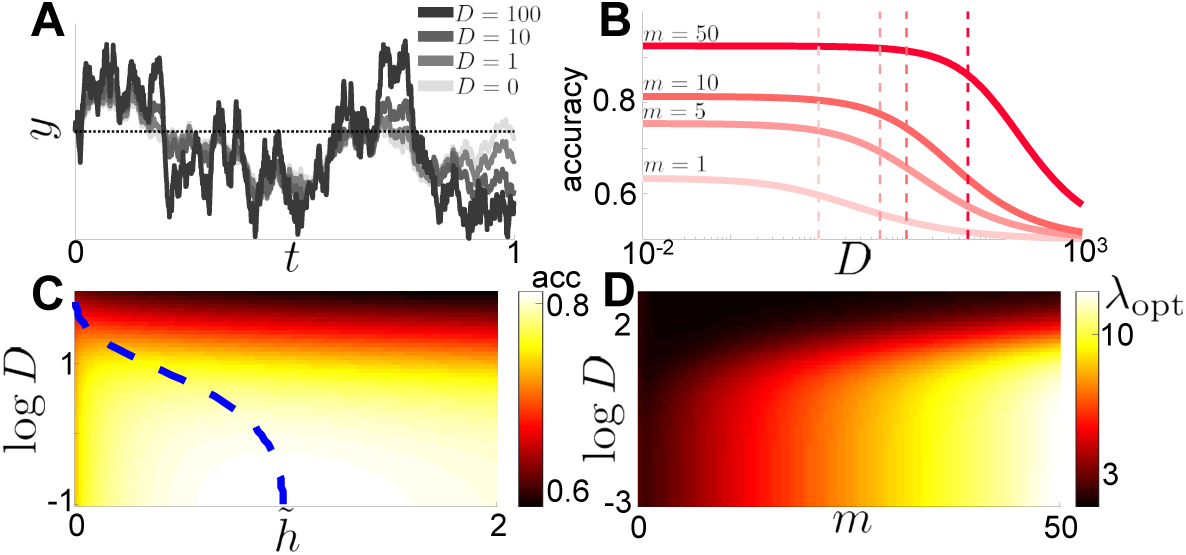
Internal noise reduces accuracy of dynamic decisions. **A:** Superimposed realizations of the nonlinear model described by Eq. (11) with added internal noise as the magnitude of internal noise, *D*, is varied (legend). **B:** Performance of nonlinear model Eq. (13) as internal noise amplitude *D* is varied for different evidence strengths *m* (legend). Vertical lines correspond to *D* = *m*. **C:** Accuracy of nonlinear model Eq. (11) with added internal noise as a function of 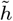 and *D* for *m* = 10. The blue line shows the optimal value of 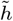 for a given *D*, when the environmental hazard rate, *h* = 1, is fixed. **D:** Leak rate *λ* that maximizes accuracy for a given evidence strength *m* and internal noise amplitude *D*.

As before, we can derive an evolution equation for the ensemble of realizations of these stochastic processes (See Appendix C). For the nonlinear Eq. (11), the corresponding differential CK equation is

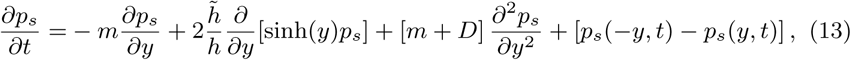

and the corresponding differential CK equation for Eq. (12) is

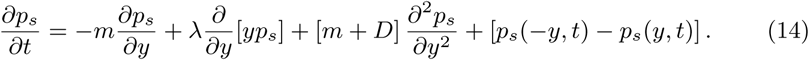

Thus, only the constants scaling the diffusion term change as compared to the models without internal noise, Eqs. (6) and (8).

How should evidence accumulation be adjusted to maximize response accuracy as the evidence strength and internal noise change? As noted earlier, humans and rats integrate internal noise in addition to information obtained from the stimulus itself (Glaze et al., 2015; Piet et al., 2018). It is therefore plausible that they adjust their stimulus integration strategies to limit the impact of internal noise on performance. Increasing the magnitude of internal noise, *D,* reduces response accuracy in the nonlinear model (Fig. 4A,B), and there there is a steep drop off in response accuracy when *D* slightly exceeds *m* (Fig. 4B). This occurs whether 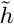 is fixed or allowed to vary. When 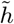 can be tuned, the value of 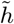 that maximizes response accuracy decreases as *D* is increased (Fig. 4C): The observer must integrate over longer timescales to average the increased internal noise and obtain a reliable estimate of the state. However, this is balanced by the need to adapt to change points as quickly as possible. Similarly, in the linear model the leak rate, *λ,* that maximizes accuracy decreases as *D* is increased (Fig. 4D). In general, as internal noise increases the observer must thus integrate over longer timescales to obtain the most accurate estimate of the state.

## 5 Discounting by bounding observer confidence

As an alternative to models that discount evidence with leak terms, we next consider models with no leak, and no-flux boundaries at *y* = ±*β* (Glaze et al., 2015). This prevents the belief from straying outside of the range −*β* ≤ *y* ≤ *β*:

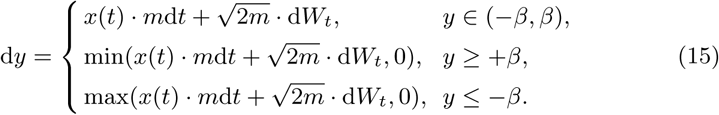

Unlike classic DDMs for two alternative free response tasks (Smith and Ratcliff, 2004; Bogacz et al., 2006; Gold and Shadlen, 2007), the process does not terminate when the belief, *y*, reaches one of the boundaries ±*β*.

More careful treatments of the reflecting boundary are possible, by considering the limit of a discrete-time biased random walk on a lattice (Erban and Chapman, 2007), but we prefer the more intuitive description of Eq. (15), whose statistics we expect to match those of more detailed models. Fig. 5A shows example realizations of this stochastic process as the boundary location, *β,* is varied, illustrating how encounters with the no-flux boundaries serve to discount evidence.

**Fig. 5.**
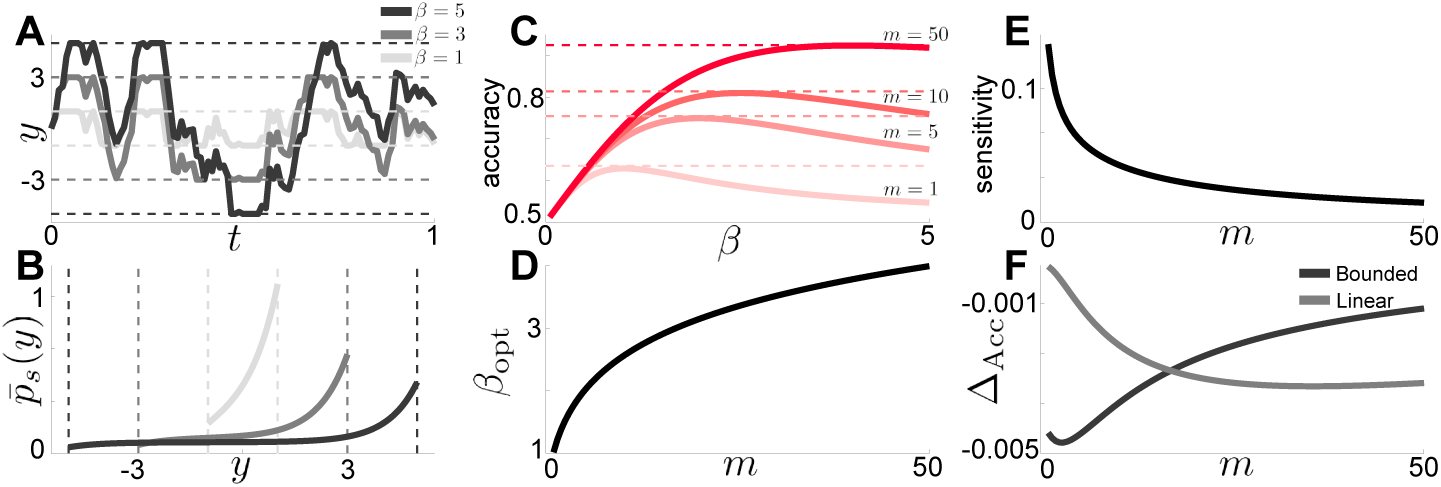
Bounded accumulator dynamics and performance. **A:** Superimposed stochastic realizations of Eq. (15) with different boundaries, *β* (legend). **B:** Superimposed steady-state distributions, 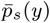, computed from Eq. (17). **C:** Tuning the boundary *β*, allows the accuracy of the bounded accumulator, Eq. (18), to closely match that of the normative model described by Eq. (2) (dashed lines) for various *m* (legend). **D:** The optimal bound, *β*_opt_, that maximizes accuracy of the bounded accumulator as a function of *m*. **E:** Sensitivity of the bounded accumulator model to changes in *β* near *β*_opt_ as a function of *m*. **F:** Difference in accuracy between the optimally tuned bounded accumulator and normative models (Δ_*Acc*_ := Acc_*B*_ (*m*) − Acc_*N*_ (*m*)) compared to accuracy difference between optimally tuned linear and normative models (Δ_*Acc*_ := Acc_*L*_(*m*) − Acc_*N*_ (*m*)) as a function of *m*.

The steady-state solution of the differential CK equations corresponding to Eq. (15) can be obtained exactly. The evolution equations are

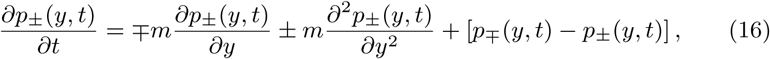

and the no flux (Robin) boundary conditions imply that

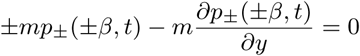

Since Eq. (16) is an advection-diffusion equation, the proper reflecting boundary is a Robin boundary (Gardiner, 2004). It can be shown that the stationary solution 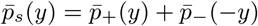 to Eq. (16) restricted by the boundary conditions is

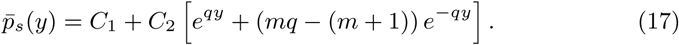

The constants *C*_1_ and *C*_2_ and details of the derivation are given in Appendix D.

The distribution is more shallow for higher *β*, as the stochastic trajectories spread over the admissible belief range (Fig. 5B). This is analogous to the sharpening (broadening) of the stationary distributions of the linear model that occurs as the leak rate is increased (decreased). Note here that *decreasing β* strengthens the discounting effect of the reflecting boundaries.

To compute steady state accuracy of the bounded accumulator model, we can integrate Eq. (17) to obtain a formula that depends on *m* and *β*:

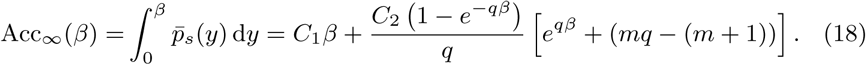

For fixed *m*, Eq. (18) varies nonmonotonically with *β*, so there is a single *β* = *β*_opt_ which maximizes the accuracy (Fig. 5C). As *m* increases, this optimal *β* increases, suggesting that as the evidence is strengthened (Fig. 5D), less discounting is needed, in contrast to the linear discounting model. Accuracy is most sensitive to changes in *β* when *m* is small (Fig. 5E).

We also compare the performance of the bounded accumulator model with that of the linear discounting model (Fig. 5F). At low *m*, linear discounting performs better than the bounded accumulator, obtaining accuracy closer to that of the normative model. The opposite is true at high *m*, in which case the bounded accumulator model performs better. This may be related to the fact that linear discounting better approximates the local dynamics of the nonlinearity −2*h* sinh(*y*) when *m* is small (and thus *y* is closer to 0), whereas a sharp boundary better approximates the strong discounting of the nonlinearity at higher values of *y* (reached when *m* is large) (Glaze et al., 2015). Both models perform quite close to the normative model when their discounting parameters are fine tuned.

Despite the bounded accumulator model’s near optimal response accuracy, it is important to note that the distributions 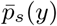 of the bounded accumulator (Fig. 5B) are very different from those of the normative (Fig. 2B) or even the linear model (Fig. 3E,F). In this respect, fitting subject confidence reports using the bounded accumulator would give very different results; we return to this point in Section 7.

## 6 Generalized discounting functions with a cubic example

There are many combinations of discounting functions and boundaries that could be used to approximate the nonlinearity 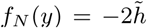 in the normative model (Wilson et al., 2010). To use our methods, we require discounting functions, *f* (*y*), that are (i) odd (*f* (−*y*) = −*f* (*y*)), and (ii) negative for some half-infinite positive region of *y* (*f* (*y*) > 0 for *y* ∈ (*a,* ∞) where *a ≥* 0); these conditions ensure convergence to non-trivial stationary distributions. For a general discounting function, the rescaled model then takes the form

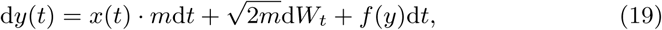

with the differential CK equation for *p*_*s*_(*y, t*) = *p*_+_(*y, t*) + *p*_−_(−*y, t*) given

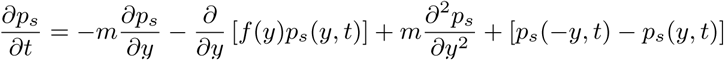

This family of evidence-discounting models could also incorporate boundary conditions as in the previous section.

A natural way to extend the linear is to introduce a cubic discounting function *f*_*C*_ (*y*) = −*λ*_1_*y* − λ_2_*y*^3^ (Piet et al., 2018) which can be tuned to better match the nonlinearity of the normative model (Fig. 6A). As shown in Section 7, this vastly improves the agreement between the stationary probability density 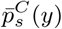 and that of the normative model, 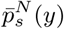, though the model is more complex (Friedman et al., 2001). Since the linear model already obtains near-optimal accuracy (Fig. 3A), we find, as expected, that the best cubic model is only slightly more accurate. In fact, accuracy drops rapidly as *λ*_2_ is changed from its optimal value (Fig. 6B). We also calculated the difference between the accuracy of the optimal cubic model and optimal linear model, Δ_Acc_(*m*) = Acc_*C*_ (*m*)−Acc_*L*_(*m*) (Fig. 6C). Accuracy improves by incorporating the cubic term at high *m* values, since nonlinear discounting is most needed at higher *y* values.

**Fig. 6.**
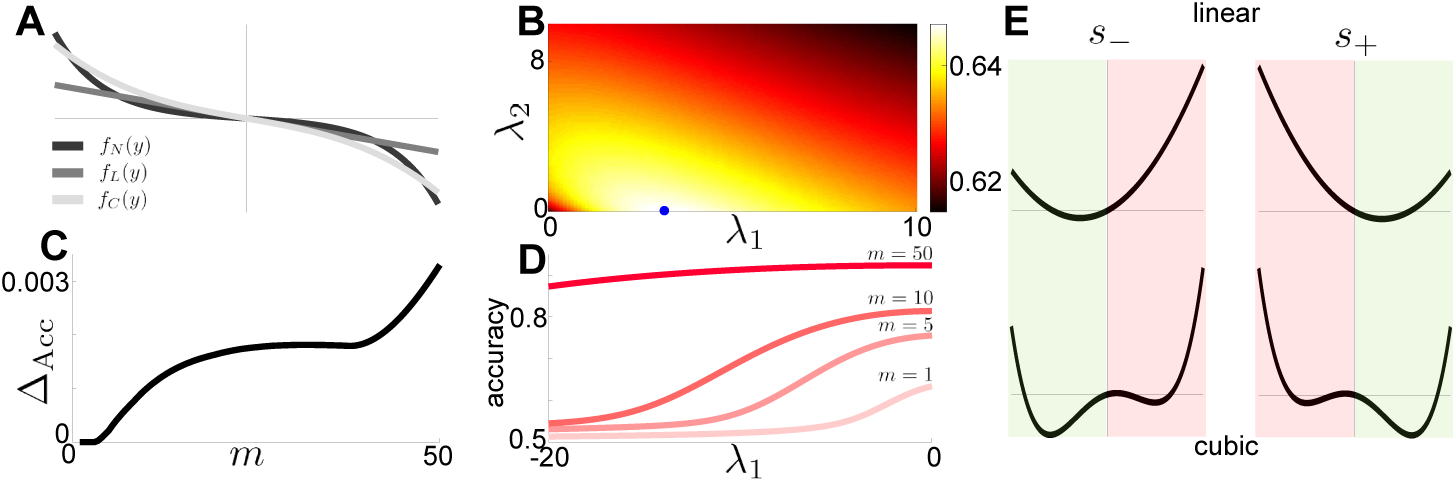
Cubic discounting functions: *f*_*C*_ (*y*) = −*λ*_1_*y* − λ_2_*y*^3^. **A:** Compared with the normative *f*_*N*_(*y*) = 2 sinh(*y*) and linear *f*_*L*_(*y*) = −*λ*_1_*y* discounting functions. **B:** Accuracy as a function of discounting rates (*λ*_1_, *λ*_2_), for *m* = 1. Dot denotes maximizer. **C:** Difference in accuracy between optimally tuned cubic and linear models (Δ_Acc_ := Acc_*C*_ (*m*) Acc_*L*_(*m*)) increase with *m*. **D:** Accuracy as a function of *λ*_1_ for different *m* (legend). **E:** Schematic of potential functions of linear model (top) and cubic model (bottom) given the state *s*(*t*). The cubic model has been mistuned so that *λ*_2_ = 1 and *λ*_1_ < 0, resulting in a double-potential well function.

However, mistuning the cubic model can considerably limit accuracy when the attractor structure of Eq. (19) with *f* (*y*) = *f*_*C*_ (*y*) is qualitatively changed (Fig. 6D). Equilibria of the noise-free model are identified by fixing *x*(*t*) = ±1 and solving the cubic equation 0 = ±*m* − λ_1_*y* − λ_2_*y*^3^. Fixing *λ*_2_ > 0, we find a critical value, 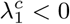 for which the attractor structure of the model switches from a single stable fixed point to two (Fig. 6E). Such bistable systems can be advantageous for working memory (Brody et al., 2003), but can hinder belief switches necessary in dynamic environments after state transitions. Fig. 6D shows that the accuracy of the model decreases as *λ*_1_ is decreased and the potential wells deepen. In these cases, the observer retains an erroneous belief long after the state has changed.

Thus, the cubic nonlinearity only marginally improves accuracy, but can have deleterious effects if mistuned. However, we may wish to use other measures of a subject’s belief to fit and validate models. In the next section we therefore ask how the full belief distribution changes with the choice of discounting function, and use KL divergence to quantify differences between different models.

## 7 Revisiting KL divergence for fitting observer belief distributions

Subject reports of confidence in decision-making tasks can be associated with LLRs of normative evidence accumulation models (Kiani and Shadlen, 2009). Thus, it may be possible to empirically estimate the belief distribution, *p*_*s*_(*y, t*), represented by our models, by asking subjects to report confidence in their choice. This provides an additional advantage of our approach over Monte Carlo simulations, as using the latter to estimate belief distributions can be costly and inaccurate (See Fig. 9 in Appendix B). As we show here, a better understanding of how our normative and approximate models deviate from one another can be gleaned by comparing their belief distributions and computing KL divergence measures.

We provide intuition for the differences we will see by plotting single stochastic realizations of all four models (Fig. 7A). The linear and (even better) the cubic models closely track the belief trajectory of the normative model, while the bounded accumulator model strays the farthest. Comparing belief distributions of all four models that minimize KL divergence with the normative model at *m* = 50 (Fig. 7B), we see that the cubic model matches the normative model far better than the linear model. This is due to the nonlinearity incorporated by the cubic term, which attenuates the tail of the distribution at high *y* values. On the other hand, the best fit bounded accumulator distribution is far from that of the normative model.

**Fig. 7.**
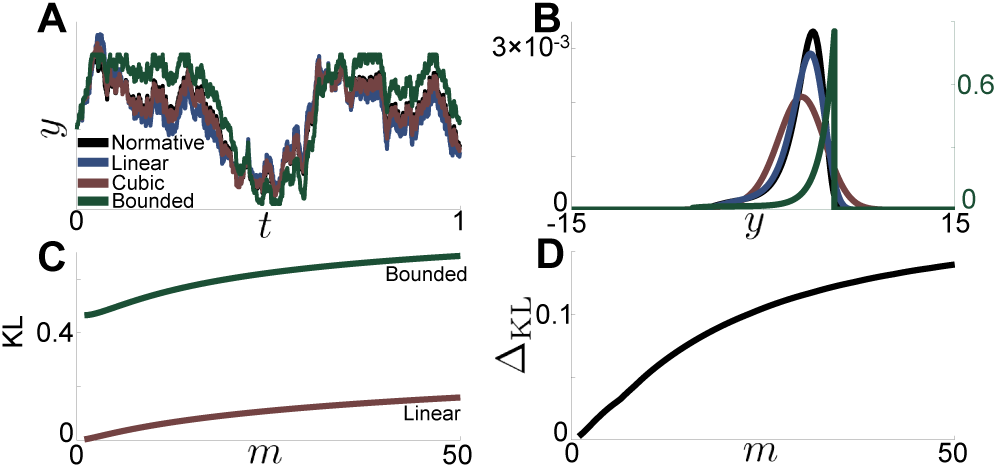
Minimizing KL divergence in our approximate models. **A:** Single realization of linear, cubic, and bounded models tuned to minimize KL divergence *m* = 50. Normative realization is shown for comparison (legend). **B:** Distributions of linear, cubic, and bounded models tuned to minimize KL divergence, again for *m* = 50. Left vertical axis gives normative, linear, and cubic scale, while right axis gives bounded accumulator scale. **C:** Minimum KL divergence for linear and bounded models as a function of *m*. **D:** Difference in KL between optimal linear and cubic models (Δ_KL_ := KL_*L*_(*m*) − KL_*C*_ (*m*)) as a function of *m*.

Computing the KL divergence between models, we arrive at two main conclusions: First, despite the fact that the bounded accumulator obtains near optimal accuracy, the corresponding belief distribution, 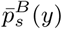, deviates from that of the normative model, 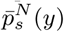, at all evidence strength values, *m* (Fig. 7C). On the other hand, though the cubic model only mildly increases response accuracy over the linear model, the corresponding belief distribution, 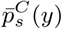, matches that of the normative model, 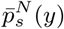, far better (Fig. 7D). Our differential CK framework allowed us to obtain these results quickly and accurately.

## 8 Chapman-Kolmogorov equations for clicks-task models

Thus far, we have been concerned with models that represent evidence accumulation in a RDMD task (Glaze et al., 2015), in which subjects receive a continuous flow of evidence during a trial. We next examine models of observers accumulating discretely timed, pulsatile evidence.

Piet et al. (2018) showed rats can perform an auditory clicks task of this type near-optimally. In this experiment, subjects are presented with two trains of clicks (one to the left ear, the other to the right), each generated by a Poisson process with instantaneous rates *r*_*L*_(*t*) and *r*_*R*_(*t*). The rates evolve according to a two-state continuous time Markov process with hazard rate *h* so that P(*r*_*j*_(*t* + d*t*)≠*r*_*j*_(*t*)) = *h*·*dt* + *o*(d*t*), *r*_*R*_(*t*)≠ *r*_*L*_(*t*) always, and *r*_*R,L*_(*t*) ∈ *r*± with *r*_+_ > r_−_. We define the state (*r*_*R*_(*t*), *r*_*L*_(*t*)) = (*r*_±_, *r*_∓_) as *s*±. Observations ξ(*t*) are now comprised of the presence or absence of left or right clicks at each time *t*. See Piet et al. (2018); Radillo et al. (2019) for details. At an interrogation time, *T*, the observer must respond which side currently has the higher click rate *r*_+_.

The model for an ideal observer’s belief 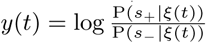 is given by

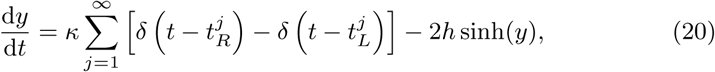

where 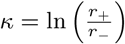 is the height of each evidence increment, 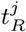 and 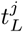 are the right and left click times, and *h* is the hazard rate. Additionally, we define the inputs’ signal-to-noise ratio (SNR) as 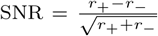 (Skellam, 1946). See Radillo et al. (2019) for a detailed discussion of how the SNR shapes model response accuracy.

Rather than carrying out another detailed analysis of several different approximations and perturbations to Eq. (20), we simply wish to show that our CK approach works and provides useful insights for click stimulus models. To sample the space of possible approximate models, we fix *h* = 1 and focus on a linear discounting and bounded accumulator model. Since Piet et al. (2018) was specifically interested in internal noise perturbed versions of the linear model, we start with this example, and then consider a bounded accumulator without internal noise that affords us explicit results.

The linear model with internal noise takes the form (Piet et al., 2018)

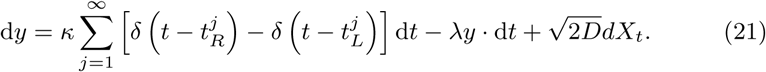

Here *λ* is the leak rate, *D* is the strength of the internal noise, and *X*_*t*_ is a Wiener process. We then define conditional densities *p*_+_(*y, t*) and *p*_−_(*y, t*) as before, writing coupled differential CK equations as

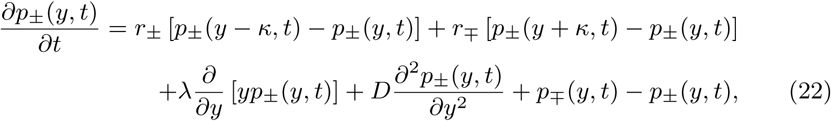

Unlike our differential CK equations for models with continuously arriving evidence, the pulses flow probability between *y* ± *κ* and *y*, preventing us from combining *p*±(*y, t*) with a change of variables and obtaining a single CK equation. Simulating Eq. (22) directly, we study how response accuracy depends on the leak and the click rates (Radillo et al., 2019). Similar to our linear model with a drift diffusion signal, accuracy varies nonmonotonically with *λ* (Fig. 8A), and is maximized at *λ* = *λ*_opt_ for a given pair (*r*_+_, *r*_−_) as plotted in Fig. 8B. Fixing SNR does not fix *λ*_opt_ as Radillo et al. (2019) showed for the normative model. As either *r*_+_ or *r*_−_ is increased, *λ*_opt_ increases (Fig. 8B), suggesting that increasing the rate of true or erroneous pulses warrants stronger evidence discounting.

**Fig. 8.**
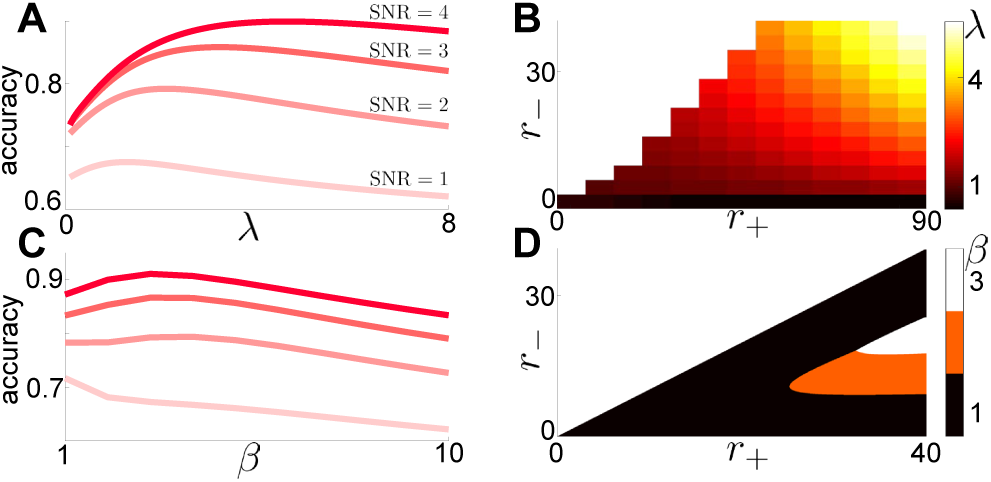
Performance of heuristic strategies on the dynamic clicks task. **A:** Accuracy of the linear model varies nonmonotonically with the leak rate, *λ* for different 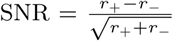 (Radillo et al., 2019) with *r*_−_ = 30 is fixed. **B:** Heatmap of optimal leak rate, *λ* = *λ*_opt_, as a function of *r*_+_ and *r*_−_ for the linear model. **C:** Accuracy of the bounded accumulator model given by Eq. (24) varies with the boundary value *β* for different SNR as *r*_−_ = 30 is fixed. **D:** Heatmap of optimal *β* = *β*_opt_ as a function of *r*_+_ and *r*_−_ for bounded accumulator clicks model.

**Fig. 9.**
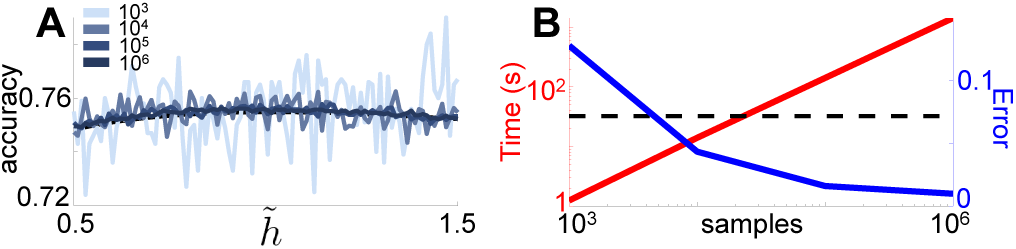
Comparison of CK equations with Monte Carlo sampling. **A:** Calculation of accuracy of mistuned nonlinear leak model for *m* = 5. Monte Carlo simulations run with varied number of samples superimposed (legend). **B:** Runtime (red) and *L*^2^ error (blue) of Monte Carlo simulations as a function of sample size. Runtime of CK equations (black dashed) superimposed for comparison. *L*^2^ error of Monte Carlo simulations calculated against results from CK equations.

We also consider a bounded accumulator with no explicit discounting function. In parallel with Eq. (16), we define the model as

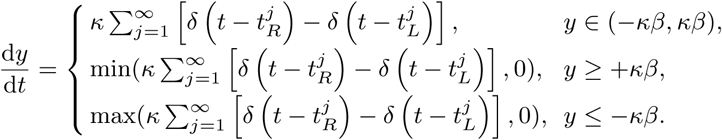

The observer’s belief is restricted to the interval [−*κβ, κβ*] for some positive integer *β* ∈ ℤ_>0_. The corresponding differential CK equations can be written as a discretized system since *y* only visits integer multiples of *κ* between −*κβ* to *κβ*. Rescaling *n* = *y/κ* yields the following system

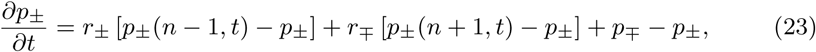

for *n* = −*β* + 1, *…, β* − 1, along with the boundary equations

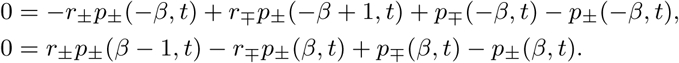

As with the continuum version of the bounded accumulator, the stationary solution 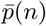 can be obtained explicitly

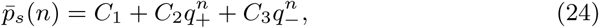

with constants *q*±, *C*_1_, *C*_2_, and *C*_3_ and the derivation given in Appendix E.

The dependence of the optimal *β* = *β*_opt_, which maximizes response accuracy, on *r*_+_ and *r* is nuanced. Fixing *r* = 30, for a given 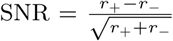, there is an optimal *β* = *β*_opt_ maximizing response accuracy (Fig. 8C). However, there is a surprisingly large range of (*r*_+_, *r*_−_) values within which *β* = 1 is optimal (Fig. 8D). There is a range of (*r*_+_, *r*_−_) for which the discounting timescale (breadth of the interval [−*βκ, βκ*]) increases with task difficulty, as long as *r*_+_ is sufficiently far from *r*_−_. However, when *r*_−_ ≈ *r*_+_, *β*_opt_ = 1, despite task difficulty. We conjecture this bound improves performance by instituting a “two click” strategy, in which the observer needs to only hear two clicks on the high click rate side to register a correct current belief. This limits the size of erroneous excursions wrong clicks can cause, as the bounds limit the effect of many misatributed clicks in a row.

Our methods thus extend to models with pulsatile evidence accumulation, illustrating their broad applicability. We can efficiently study model performance and its dependence on task parameters, and even explicitly analyze the resulting equations to determine how approximate models perform.

## 9 Conclusion

Decision-making models are key to understanding how animals integrate evidence to make choices in nature. Animals most likely use heuristic strategies in dynamic tasks as they can be easier to implement, and have utility that is close to optimal (Rahnev and Denison, 2018). Normative models are still useful, however, as subject performance can be benchmarked against them, allowing possible insights into how and why organisms fail to perform optimally (Geisler, 2003). Investigating optimal models and their approximations requires simulations across large parameter spaces; these necessarily require rapid simulation techniques to obtain refined results. Efficient computational methods are therefore essential for the analysis of evidence accumulation models, and their application to experiment design.

Using differential CK equations to describe ensembles of decision model realizations speeds up computation and describes the time-dependent probability density of an observer’s belief. Thus, traditional metrics of performance (e.g., accuracy) and other less common model comparison metrics (KL divergence) can be computed rapidly. This opens new avenues for comparing normative and heuristic decision making models, and for determining task parameter ranges to distinguish models. There is also hope that in high throughput experiments, sufficient data could be collected to specify subject confidence distributions, which could be fit, or compared to model predictions (Piet et al., 2019).

Doubly stochastic and jump-diffusion models appear in a number of other contexts in neuroscience and beyond (Hanson, 2007; Horsthemke and Lefever, 2006). For instance, dichotomous and white noise have been included in linear integrate and fire (LIF) models to model voltage or channel fluctuations (Droste and Lindner, 2014, 2017; Salinas and Sejnowski, 2002). The interspike interval statistics of these models can be analyzed directly by considering the corresponding differential CK equations. Unlike the models we consider here, the LIF model includes a single absorbing boundary and reset condition, which must be treated carefully when defining the flow of probability through state space.

We have studied a number heuristic models and computed how their performance depends on both task parameters and evidence discounting parameters. Well tuned heuristic models, such as the linear and bounded accumulator models, can in fact exhibit near-normative performance (Glaze et al., 2015; Veliz-Cuba et al., 2016; Radillo et al., 2019). There are specific parameter regimes (low versus high *m*) in which certain heuristic models perform better; our differential CK methods have allowed us to explore these regimes rapidly. Importantly, Brunton et al. (2013), and Piet et al. (2018) have shown that internal noise determines subject performance in decision tasks, in addition to the variability of the signal. We have confirmed that including internal noise causes optimal evidence-discounting to be weakened as noise increases, and that accuracy drops off precipitously once the amplitude of internal noise reaches that of the signal.

Our approach can also be used by experimentalists testing observer performance in dynamic-decision tasks. Models can thus guide one’s choice of task parameters when setting up experiments to determine the strategies subjects use to make decisions in dynamic environments. As in Radillo et al. (2019), we found that accuracy is most sensitive to one’s choice of model and tuning when tasks are of intermediate difficulty. In contrast, tasks that are easy (hard) are performed well (poorly) by most models. Also, the full belief distributions generated by our methods could be subsampled to produce randomized responses for comparison with subject data (Drugowitsch, 2016). It may also be feasible to use our differential CK equations to model trial-to-trial belief distributions of subjects, as affected by internal noise hidden to the experimentalist. This approach was recently developed in Piet et al. (2019) to account for for variability in subject responses.

Model development can also help to inspire new experimental tasks, based on predictions and ideas that arise from mathematically describing subjects’ decision processes. One possible extension of the tasks we have discussed here could consider stochastic switches in evidence quality within trials. Past work has focused on both theoretical predictions and experimental results associated with task difficulty switching between trials (Drugowitsch et al., 2012; Zhang et al., 2014), suggesting subjects’ decision thresholds may vary with time as task difficulty is inferred throughout the trial. When both the state and difficulty switch stochastically within a trial, the effective state is governed by a multi-state continuous time Markov process. Details that could be introduced into such multi-state models, such as asymmetric evidence qualities and the ability to turn off evidence, offer a rich framework for applying the stochastic methods we have developed here. Another extension we could consider in our models is the recently developed click task model with stochastically drawn click heights (Piet et al., 2018; Radillo et al., 2019). Jumps would then be represented by an integral over the entire belief space, requiring new computational methods for efficient simulation of the associated differential CK equations.

In recent years, decision-making models and experiments have been developed to incorporate more naturalistic scenarios in which the environment changes in fluid yet predictable ways. The associated normative models can be complex, and efficient simulation techniques are important for evaluating performance across different models and interpreting experimental decision data from psychophysics tasks. It is also important to develop families of plausible heuristic models that subjects may be implementing, and to find ways to compare them with normative models. Our Chapman-Kolmogorov framework provides a straightforward and robust way to achieve these goals.

## Code availability

Refer to https://github.com/nwbarendregt/DynamicDecisionCKEquations for the MATLAB finite difference code used to perform the analysis and generate figures.

## A Normative evidence-accumulation in dynamic environments

Here we derive the continuum limit of the Bayesian update equation for continuous evidence accumulation in a changing environment. Starting with the discrete time model, we define *L*_*n,*±_ = P(*s*(*t*_*n*_) = *s*±|ξ_1:*n*_) as the probability of being in state *s*± at time *t*_*n*_ assuming a sequence of observations ξ_1:*n*_. The state *s*(*t*) changes between evenly spaced time points *t*_1:*n*_ (with Δ*t* := *t*_*n*_ - *t*_*n-*1_) at a hazard rate *h*_Δ*t*_ := *h* · Δ*t* := P(*s*(*t*_*n*_) = *s*∓|*s*(*t*_*n-*1_) = *s*±). The likelihood function *f*_Δ*t*,±_(ξ) = P(ξ|*s* = *s*±; Δ*t*) is the conditional probability of observing sample ξ given state *s*±, parameterized by Δ*t*.

We begin by assuming an ideal observer who knows the environmental hazard rate *h*. Using Bayes’ rule and the law of total probability, we can relate *L*_*n*,±_ to the probability at the previous time step according to the weighted sum (Veliz-Cuba et al., 2016)

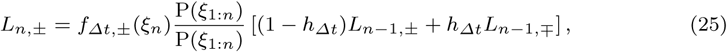

where *L*_0,±_ = P(*s*(*t*_0_) = *s*±). Defining 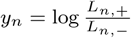, we can compute

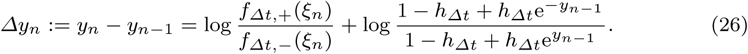

In search of the continuum limit of this equation, we assume 0 < Δ*t*≪ 1, 0 < |Δ*y*_*n*_| ≪1, and use the approximation log(1 + *z*) ≈ *z* to obtain

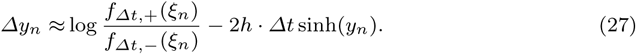

Replacing the index *n* with the time *t* and applying the functional central limit theorem as in Billingsley (2008); Bogacz et al. (2006), we can write Eq. (27) as

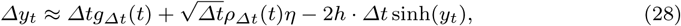

where *η* is a random variable with a standard normal distribution and

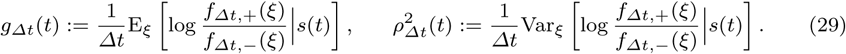

The drift *g*_Δ*t*_ and variance 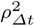 diverge unless *f*_Δ*t,*±_(ξ) are scaled appropriately in the Δ*t* → 0 limit. A reasonable assumption that can be made to compute *g*_Δ*t*_ and 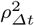 explicitly is to take observations ξ to follow normal distributions with mean and variance scaled by Δ*t* (Bogacz et al., 2006; Veliz-Cuba et al., 2016)

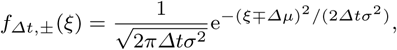

so we can compute the limits of Eq. (29) as

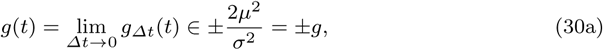

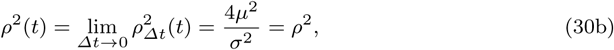

where *g*(*t*) ∈ {+*g*, −*g*} is a telegraph process with probability masses P(±*g, t*) evolving as 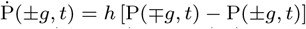 and *ρ*^2^(*t*) = *ρ*^2^ remains constant. Therefore, the continuum limit (Δ*t* → 0) of Eq. (28) is

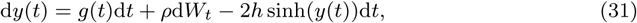

where d*W* is a standard Wiener process. Eq. (31) provide the normative model of evidence accumulation for an observer who knows the hazard rate *h* and wishes to infer the sign of *g*(*t*) at time *t* with maximal accuracy (Glaze et al., 2015; Veliz-Cuba et al., 2016).

However, we are also interested in near-normative models in which the observer assumes an incorrect hazard rate 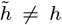. In such a case, the analysis proceeds as before, with the probabilistic inference process simply involving 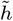 now rather than *h*, and the result is

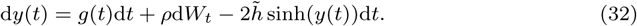

Lastly, note that if indeed the original observations ξ are drawn from normal distributions, Eq. (30) states *g*(*t*) ∈ ±*g* where *g* = 2*µ*^2^*/σ*^2^ and *ρ*^2^ = 2*g*. Rescaling time *ht*↦*t*, we can then express Eq. (32) in terms of the following rescaled equation

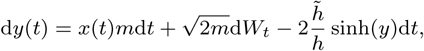

where *m* = 2*µ*^2^*/*(*hσ*^2^) and *x*(*t*) ∈ ±1 is a telegraph process with hazard rate 1, as shown in Eq. (2) of the main text.

## B Finite difference methods for Chapman-Kolmogorov equations

We use a finite difference method to simulate the differential CK equations. The method is exemplified here for the normative CK equation from Eq. (6), but a similar approach is also used for the linear, cubic, and pulsatile equations. For stability purposes, our method uses centered differences in *y* and backward-Euler in *t*. This gives the following finite difference approximations of the functions and their derivatives in Eq. (6):

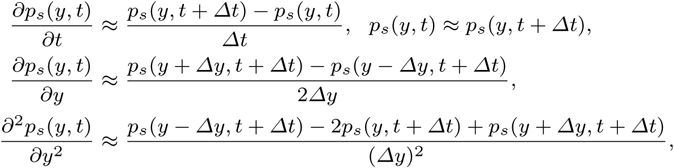

where Δ*t* and Δ*y* are timestep and spacestep of the simulation, respectively. Substituting into Eq. (6) and solving for *p*_*s*_(*y, t*) at each point on a mesh **y** for *y* gives the system of equations:

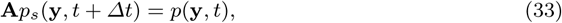

where **A** is tridiagonal with elements along the primary off-diagonal. This system can be inverted at each timestep and used to calculate the updates *p*_*s*_(**y**, *t* + Δ*t*).

For the boundary conditions, we impose no-flux conditions at the mesh boundaries ±*b*. For a standard drift-diffusion equation with drift *A*(*y*) and diffusion constant *B*(*y*), this condition takes the form

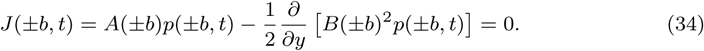

Using the finite difference approximations

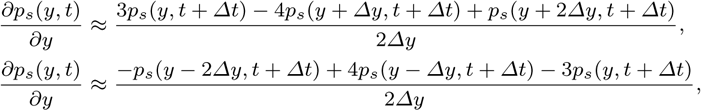

we can plug in ±*b* to the appropriate replacement and use Eq. (34) to find the appropriate boundary terms for the system in Eq. (33).

Fig. 9 shows the results of Monte Carlo simulations compared against those from the CK equations; Monte Carlo simulations are less smooth (Fig. 9A), making optimality calculations less accurate. Furthermore, obtaining results that are close to those from the CK equations takes much longer to run (Fig. 9B).

## C Deriving differential CK Equation with internal noise

Here we provide intuition for the form of the diffusion coefficient in Eq. (13) for the belief distribution of a normative observer strategy with additional internal noise of strength *D*.

Starting with the SDE in Eq. (12), because 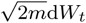 and 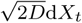 are increments of independent Wiener processes, we can define a new Wiener process 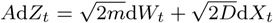 that has the same statistics as the original summed Wiener processes (Gardiner, 2004). To determine the appropriate effective diffusion constant *A*, we note that

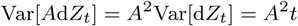

and

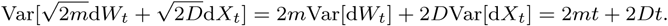

This requires 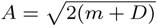, and means Eq. (12) can be rewritten as

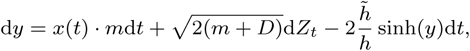

which following Gardiner (2004), has the differential CK equation given by Eq. (13).

## D Steady state solution of the bounded accumulator model

Steady state solutions of Eq. (16) are derived first by noting that *∂*_*t*_*p*^±^ = 0 implies

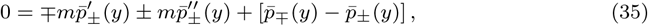

with boundary conditions 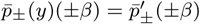. Eq. (35) has solutions 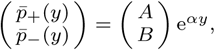, with characteristic equation *m*^2^*α*^4^–(*m*^2^ + 2*m*)*α*^2^ = 0. The characteristic roots are *α* = 0,± *q*, where we define 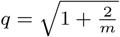. For *α* = 0, we have *A* = *B*, whereas for *α* = ±*q*, the symmetry 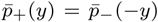 implies *B* = (*mq* − (*m* + 1))*A* for *α* = +*q* and *A* = (*mq* − (*m* + 1))*B* for *α* = *-q*. Lastly, defining the sum 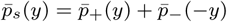, we obtain

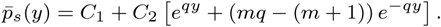

The no flux boundary conditions 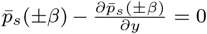 along with the normalization requirement 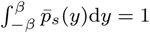 give explicit expressions for the constants

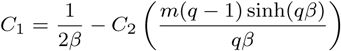

and

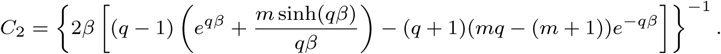

## E Steady state solution of the clicks-task bounded accumulator model

Considering Eq. (23), we look for stationary solutions of the form 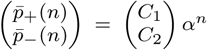, yielding the characteristic equation

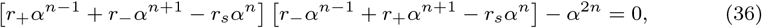

where *r*_*s*_ = *r*_+_ + *r*_−_ + 1. Solving Eq. (36) gives *α* = 1 with eigenfunction *C*_1_ = *C*_2_ and two roots *α* = *q*± of the quadratic 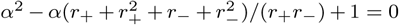. Superimposing the eigenfunctions, redefining constants, and defining 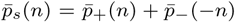 gives the general solution

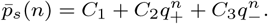

The constants *C*_1_, *C*_2_, and *C*_3_ can be determined by normalization 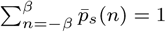 and the stationary boundary conditions

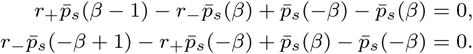

Long term accuracy of the bounded accumulator is then determined by the weighted sum 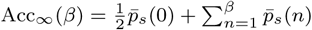.

## Acknowledgements

This work was supported by and NSF/NIH CRCNS grant (R01MH115557) and NSF (DMS-1517629). ZPK was also supported by NSF (DMS-1615737). KJ was also supported by NSF (DBI-1707400). We thank Sam Isaacson and Jay Newby for feedback on setting up the boundary value problem for the bounded accumulator model. We are also grateful to Adrian Radillo and Tahra Eissa for comments on a draft version of the manuscript.

## References

Bankó ÉM, Gál V, Körtvèlyes J, Kovács G, Vidnyánszky Z (2011) Dissociating the effect of noise on sensory processing and overall decision difficulty. Journal of Neuroscience 31(7):2663–2674

Behrens TE, Woolrich MW, Walton ME, Rushworth MF (2007) Learning the value of information in an uncertain world. Nature neuroscience 10(9):1214

Billingsley P (2008) Probability and measure. John Wiley & Sons

Bogacz R, Brown E, Moehlis J, Holmes P, Cohen JD (2006) The physics of optimal decision making: a formal analysis of models of performance in two-alternative forced-choice tasks. Psychological review 113(4):700

Brea J, Urbanczik R, Senn W (2014) A Normative Theory of Forgetting: Lessons from the Fruit Fly. PLoS Computational Biology 10(6):e1003640

Brody CD, Romo R, Kepecs A (2003) Basic mechanisms for graded persistent activity: discrete attractors, continuous attractors, and dynamic representations. Current opinion in neurobiology 13(2):204–211

Brunton BW, Botvinick MM, Brody CD (2013) Rats and humans can optimally accumulate evidence for decision-making. Science 340(6128):95–98

Busemeyer JR, Townsend JT (1992) Fundamental derivations from decision field theory. Mathematical Social Sciences 23(3):255–282

Droste F, Lindner B (2014) Integrate-and-fire neurons driven by asymmetric dichotomous noise. Biological cybernetics 108(6):825–843

Droste F, Lindner B (2017) Exact results for power spectrum and susceptibility of a leaky integrate-and-fire neuron with two-state noise. Physical Review E 95(1):012411

Drugowitsch J (2016) Fast and accurate monte carlo sampling of first-passage times from wiener diffusion models. Scientific reports 6:20490

Drugowitsch J, Moreno-Bote R, Churchland AK, Shadlen MN, Pouget A (2012) The cost of accumulating evidence in perceptual decision making. Journal of Neuroscience 32(11):3612–3628

Erban R, Chapman SJ (2007) Reactive boundary conditions for stochastic simulations of reaction–diffusion processes. Physical Biology 4(1):16

Faisal AA, Selen LP, Wolpert DM (2008) Noise in the nervous system. Nature reviews neuroscience 9(4):292

Friedman J, Hastie T, Tibshirani R (2001) The elements of statistical learning, vol 1, Springer series in statistics New York, NY, USA:, chap 7: Model Assessment and Selection

Gardiner C (2004) Handbook of stochastic methods: for physics, chemistry & the natural sciences,(series in synergetics, vol. 13)

Geisler WS (2003) Ideal observer analysis. The visual neurosciences 10(7):12–12

Glaze CM, Kable JW, Gold JI (2015) Normative evidence accumulation in unpredictable environments. Elife 4:e08825

Glaze CM, Filipowicz AL, Kable JW, Balasubramanian V, Gold JI (2018) A bias–variance trade-off governs individual differences in on-line learning in an unpredictable environment. Nature Human Behaviour 2(3):213

Gold JI, Shadlen MN (2007) The neural basis of decision making. Annual review of neuroscience 30

Hanson FB (2007) Applied stochastic processes and control for Jump-diffusions: modeling, analysis, and computation, vol 13. Siam

Horsthemke W, Lefever R (2006) Noise-Induced Transitions: Theory and Applications in Physics, Chemistry, and Biology. Springer Series in Synergetics, Springer Berlin Heidelberg

Kiani R, Shadlen MN (2009) Representation of confidence associated with a decision by neurons in the parietal cortex. science 324(5928):759–764

Moehlis J, Brown E, Bogacz R, Holmes P, Cohen JD (2004) Optimizing reward rate in two alternative choice tasks: Mathematical formalism. Center for the Study of Brain, Mind and Behavior, Princeton University pp 04–01

Ossmy O, Moran R, Pfeffer T, Tsetsos K, Usher M, Donner TH (2013) The timescale of perceptual evidence integration can be adapted to the environment. Current Biology 23(11):981– 986

Piet A, Hady AE, Boyd-Meredith T, Brody C (2019) Neural dynamics during changes of mind. In: Computational and Systems Neuroscience 2019 Lisbon, Portugal

Piet AT, El Hady A, Brody CD (2018) Rats adopt the optimal timescale for evidence integration in a dynamic environment. Nature communications 9(1):4265

Radillo AE, Veliz-Cuba A, Josić K, Kilpatrick ZP (2017) Evidence accumulation and change rate inference in dynamic environments. Neural computation 29(6):1561–1610

Radillo AE, Veliz-Cuba A, Josić K (2019) Performance of normative and approximate evidence accumulation on the dynamic clicks task. Neurons, Behavior, Data analysis, and Theory submitted

Rahnev D, Denison RN (2018) Suboptimality in perceptual decision making. Behavioral and Brain Sciences 41:e223, DOI 10.1017/S0140525X18000936

Ratcliff R (1978) A theory of memory retrieval. Psychological review 85(2):59

Ratcliff R, McKoon G (2008) The diffusion decision model: theory and data for two-choice decision tasks. Neural computation 20(4):873–922

Salinas E, Sejnowski TJ (2002) Integrate-and-fire neurons driven by correlated stochastic input. Neural computation 14(9):2111–2155

Skellam JG (1946) The frequency distribution of the difference between two poisson variates belonging to different populations. Journal of the Royal Statistical Society Series A (General) 109(Pt 3):296–296

Smith PL (2010) From poisson shot noise to the integrated ornstein–uhlenbeck process: Neurally principled models of information accumulation in decision-making and response time. Journal of Mathematical Psychology 54(2):266–283

Smith PL, Ratcliff R (2004) Psychology and neurobiology of simple decisions. Trends in neurosciences 27(3):161–168

Urai AE, Braun A, Donner TH (2017) Pupil-linked arousal is driven by decision uncertainty and alters serial choice bias. Nature Communications 8:14637

Van Den Berg R, Anandalingam K, Zylberberg A, Kiani R, Shadlen MN, Wolpert DM (2016) A common mechanism underlies changes of mind about decisions and confidence. Elife 5:e12192

Veliz-Cuba A, Kilpatrick ZP, Josic K (2016) Stochastic models of evidence accumulation in changing environments. SIAM Review 58(2):264–289

Wilson RC, Nassar MR, Gold JI (2010) Bayesian online learning of the hazard rate in change-point problems. Neural computation 22(9):2452–2476

Yu AJ, Cohen JD (2008) Sequential effects: Superstition or rational behavior? Advances in Neural Information Processing Systems 21:1873–1880

Zhang S, Lee MD, Vandekerckhove J, Maris G, Wagenmakers EJ (2014) Time-varying boundaries for diffusion models of decision making and response time. Frontiers in Psychology 5:1364

